# Generalizing the Gaussian Network Model: Spanning-Tree Thermodynamics Shows Entropy-Driven KRAS Activation

**DOI:** 10.64898/2026.02.25.707891

**Authors:** Fatma Senguler Ciftci, Burak Erman

## Abstract

The GTPase KRAS executes a conformational switch between a GTP-bound active state and a GDP-bound inactive state, a process central to oncogenic signalling. However, the structural basis of this switching at the level of residue-contact organization remains incompletely characterized by traditional binary structural models. Here, we present a statistical-mechanical generalization of the Gaussian Network Model (GNM) by constructing spanning-tree partition functions for residue-contact graphs using the weighted Kirchhoff Laplacian in conjunction with the Matrix-Tree Theorem.

Within this framework, the standard GNM is recovered in the high-temperature limit, whereas the present formulation enables a continuous Boltzmann-weighted ensemble analysis. We compute the network free energy *F*, mean contact energy *Ē*, heat capacity *C*_*v*_, and thermodynamic entropy *S* across an effective temperature sweep that maps the combinatorial diversity of the contact network, thereby probing the topological landscape rather than structural melting.

Differential analysis reveals that KRAS activation reflects a systematic entropy-enthalpy compensation mechanism: the active state incurs a systematic energetic penalty (Δ*Ē* > 0) that is offset by a marked gain in conformational entropy (Δ*S* > 0), with a free-energy crossover occurring at *kT* ≈ 2.41 Å. Edge marginal inclusion probabilities, obtained via effective-resistance theory, identify Switch I (residues 25-40) as the primary allosteric locus of nucleotide-driven network reorganization.

This approach provides a thermodynamically grounded perspective on KRAS allostery, quantitatively demonstrating how network architecture enables functional versatility through entropy-driven conformational flexibility.

**Entropy-driven KRAS activation revealed by spanning-tree thermodynamics:** Spanning-tree partition functions built from weighted residue-contact graphs show that KRAS activation involves entropy-enthalpy compensation. The GDP-bound network (left) is dominated by a small number of low-energy spanning trees, whereas the GTP-bound network (right) accesses a larger ensemble with greater topological diversity. The active state pays an energetic cost (Δ*Ē* > 0) offset by a conformational entropy gain (Δ*S* > 0), with Switch I (residues 25-40, dashed box) emerging as the primary site of nucleotide-driven network reorganisation.

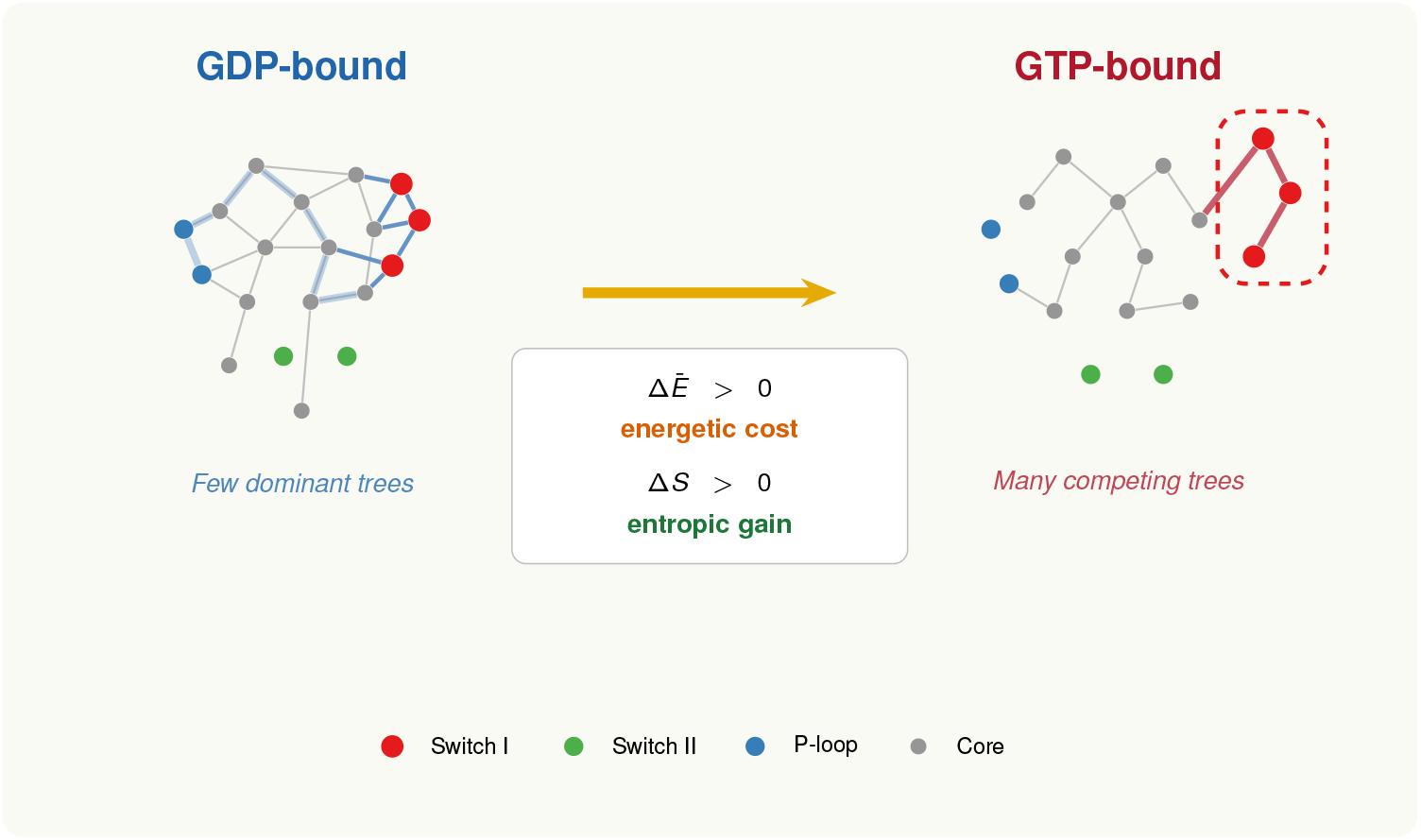

## 1 INTRODUCTION

KRAS is the most frequently mutated oncogene in human cancer: activating mutations at positions G12 and G13 appear in roughly 30% of all tumours. ^1^ Functioning as a molecular switch, the protein cycles between GTP-bound active and GDP-bound inactive conformations to regulate proliferation, survival, and differentiation through downstream effectors including RAF, PI3K, and RALGDS. ^2^ This clinical importance has driven intensive efforts toward direct small-molecule inhibitors. ^3^ Thermodynamic and kinetic aspects of KRAS allostery and its downstream interactions have been studied by Manley and Lin, ^4^, while Killoran and Smith ^5^ have characterised nucleotide cycling and effector interactions across multiple GTPases at conformational resolution. A substantial and growing body of work addresses KRAS allostery through GTP-dependent conformational dynamics, reorganisation of interaction networks, and thermodynamic analyses of binding energetics. ^6–8^ Integrative approaches combining Markov state modelling with allosteric network analysis have further mapped KRAS-effector binding landscapes and shown how mutations redistribute communication pathways. ^9^ These studies have mapped allosteric communication through conformational sampling, mutational scanning, and network centrality analysis, yet the thermodynamic consequences of global contact-network reorganisation upon nucleotide exchange have not been assessed quantitatively. A central open question is how the global architecture of residue contacts, rather than any single interaction, collectively enables or restricts allosteric signal transmission between the nucleotide-binding pocket and distal effector surfaces. This question is especially acute for KRAS, where global network reorganisation upon nucleotide exchange may be a more fundamental determinant of functional switching than local structural rearrangements alone.

Conventional structural analyses compare active and inactive states by cataloguing residue pairs that gain or lose contacts upon nucleotide exchange. ^10,11^ Elastic network models extend this picture with coarse-grained descriptions of collective motions. Still, because they are defined about a single energy minimum, they cannot capture the multiplicity of communication pathways accessible to the network. ^12^ Recent work has advanced the field by deriving edge centrality measures that implicitly enumerate communication pathways through the matrix-tree theorem, revealing conserved allosteric bottlenecks in KRAS. ^13^ That framework, however, does not assign explicit statistical weights to individual spanning-tree configurations and therefore cannot yield a temperature-dependent partition function from which heat capacity, switching free energy, and entropic contributions to state transitions follow. Neither class of approach provides what is needed: a thermodynamically consistent partition function that assigns statistical weights to contact configurations and yields well-defined free energy, entropy, and heat capacity for the network as a whole. Without such a description, the thermodynamic cost of switching between states and the relative contributions of energetic versus entropic terms cannot be assessed from static structural data. The framework introduced here addresses this gap directly by summing over the complete ensemble of contact network configurations, yielding thermodynamic quantities that are immediately comparable between the two nucleotide states.

Protein contact networks exhibit small-world communication properties, with short path lengths that enable efficient signal propagation across the structure. ^14^ We represent the protein’s residue-contact network as a spanning tree, an acyclic subgraph that includes all N residues and connects them using exactly N - 1 contacts. Each spanning tree constitutes a minimal allosteric backbone: the smallest subset of contacts through which a perturbation at the nucleotide-binding pocket can reach every other residue without redundancy. The partition function enumerates all such backbones, each weighted by its Boltzmann factor, and thereby quantifies the combinatorial diversity of allosteric routes available at a given effective temperature. Earlier approaches have characterised network topology through weighted Laplacian spectra and perturbation analysis, ^15^ or through centrality rankings and coevolutionary methods. ^16^ The present work extends this programme by assigning explicit Boltzmann weights to individual spanning-tree configurations, converting the spectral description into a statistical-mechanical ensemble from which thermodynamic potentials follow exactly.

Our formulation is grounded in algebraic graph theory. ^17,18^ Each KRAS structure is represented as a weighted residue contact graph, and its spanning-tree partition function is evaluated as a Boltzmann-weighted sum over all connected, cycle-free subgraphs that span the full residue set. This statistical-mechanical description shows how allosteric signals distribute across competing pathways, with each tree configuration’s weight reflecting its total interaction energy at a given effective temperature. Changes in this combinatorial diversity upon nucleotide exchange are invisible to pairwise contact analyses or single-minimum elastic descriptions; it is precisely these changes that, we argue, define the thermodynamic distinction between the active and inactive states.

A related graph-theoretic approach was developed by Senet and co-workers, who defined a local entropy based on subgraph enumeration around each residue to quantify conformational heterogeneity across folded, unfolded, and intrinsically disordered states. ^19^ Their local entropy functions as an order parameter for protein unfolding and correlates with NMR chemical shifts. The present work takes a global perspective: the partition function sums over all connected, cycle-free subgraphs spanning the entire residue set, yielding network-level free energy, entropy, and heat capacity rather than residue-level disorder measures.

The theoretical foundation of the Gaussian Network Model (GNM) derives from the theory of elastic network constraints established by Flory and Erman ^22^, who established the early physical basis for modeling junction fluctuations in constrained systems. The model ^23^ approximates residue interactions with a binary contact function: residues within a cutoff distance interact with uniform strength; those beyond do not interact at all. Computationally convenient, this replaces the continuous spectrum of interaction strengths with a present/absent dichotomy. We adopt a different approximation. A hard cutoff *d*_*c*_ = 8.0 Å defines which residue pairs form edges in the contact graph; within this graph, each edge receives a continuous Boltzmann weight that decays smoothly with inter-residue distance. This replaces the uniform-strength assumption of the GNM with a heterogeneous weighting that transforms the network from a single-minimum harmonic system into a statistical-mechanical ensemble over all connected configurations. This transition from a purely topological representation of a continuously weighted interaction landscape is what enables the derivation of thermodynamic potentials from the elastic framework. As shown formally in § 2.8, the standard GNM is recovered as the high-temperature limit of this ensemble, where all weights converge to unity, and the partition function reduces to the spanning-tree count. The generalisation opens access to the full thermodynamic apparatus of partition functions, free energies, and entropies, all derivable exactly from network topology without additional approximation or sampling.

Applying this framework to two crystallographic wild-type KRAS structures, 6GOD (GTP-analogue-bound, active) and 4OBE (GDP-bound, inactive), we find that activation cannot be described as a purely energetic transition. Rather, it reflects entropy-enthalpy compensation: the active state incurs a systematic energetic penalty (Δ*Ē* > 0) offset by a gain in conformational entropy (Δ*S* > 0), with the free-energy balance crossing at a well-defined intrinsic temperature. Switch I (residues 25-40) emerges as the primary site of nucleotide-driven network reorganisation.

## 2 METHODS

The analysis proceeds in a sequence of deterministic steps from atomic coordinates to thermodynamic observables. No stochastic sampling, molecular dynamics, or normal-mode approximation is involved at any stage. Every parameter choice is collected in Table 3 at the end of this section; the complete source code is publicly available (§ 6).

### 2.1 Structural Data

Two wild-type KRAS crystal structures were obtained from the RCSB Protein Data Bank: **6GOD** (GTP-analogue-bound, chain A, 1.90 Å resolution, 172 resolved residues) and **4OBE** (GDP-bound, chain A, 1.60 Å resolution, 169 resolved residues). Only the C*α* atom of each residue was retained. No alignment, superposition, or renumbering was applied; residue indices match the deposited PDB numbering throughout. The two structures share identical numbering over their common range; 4OBE simply lacks Met170, Ser171, and Lys172 at the C-terminus.

Each structure thus yields an ordered list of *n* residues with three-dimensional coordinates **r**_*i*_ = (*x*_*i*_, *y*_*i*_, *z*_*i*_), where *n* = 172 for 6GOD and *n* = 169 for 4OBE.

### 2.2 ContacXt Graph and Cutoff

From these coordinates, we build a residue contact graph *G* = (*V, e*), with one node per residue (|*V* | = *n*). An undirected edge *e* = (*i, j*) is placed whenever the C*α*-C*α* distance falls below a hard cutoff:

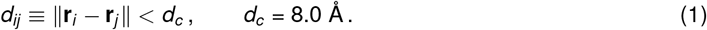

For every admitted edge, the *exact* distance *d*_*ij*_ is stored; it is not rounded to *d*_*c*_. Edge lengths range from about 3.8 Å for consecutive backbone pairs to just under 8.0 Å for the most peripheral contacts. The sensitivity of the meaningful temperature window and the contact count to the cutoff *d*_*c*_ is examined in Figure S3; the qualitative conclusions remain unchanged for all cutoffs tested. Pairs with *d*_*ij*_ ≥ *d*_*c*_ are excluded entirely (*w*_*ij*_ = 0) and play no further role. Both structures remain fully connected at *d*_*c*_ = 8.0 Å, so no residues were discarded.

The resulting networks contain 172 nodes / 830 edges (6GOD) and 169 nodes / 809 edges (4OBE). The modest size difference reflects the three extra C-terminal residues (Met170, Ser171, Lys172) resolved in the active structure; these additional nodes introduce not only themselves but also the contacts they form with neighbouring residues, accounting for the 21-edge surplus in 6GOD.

### 2.3 Edge Energies, Boltzmann Weights, and the Role of Temperature

Each edge is assigned an energy proportional to its *Cα* − *Cα* distance, which we calibrate via the effective temperature *kT*.

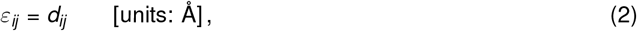

the simplest monotonic function of distance. Short contacts are energetically favoured; peripheral ones are penalised. For 6GOD the range is *ε*_min_ ≈ 3.78 Å to *ε*_max_ ≈ 7.99 Å, with a mean of roughly 5.8 Å. Robustness to alternative functional forms (quadratic, inverse, harmonic) was verified but is not reported.

At effective temperature *kT* (units: Å; see below), each edge receives a Boltzmann weight

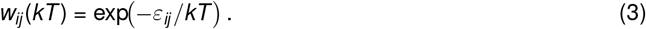

Since edge energies differ, the weights are heterogeneous at any fixed *kT* . The ratio between two edges,

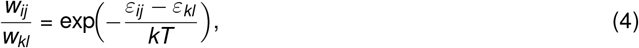

changes with temperature whenever *ε*_*ij*_ ≠*ε*_*kl*_ . The same principle extends to entire spanning trees: given two trees *τ*_1_ and *τ*_2_ with total energies *E* (*τ*_1_) and *E* (*τ*_2_) (defined in § 2.5), their probability ratio is

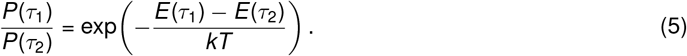

Temperature, therefore, acts as a selective force that actively redistributes statistical weight across the spanning-tree ensemble. At low *kT*, the partition function is dominated by low-energy trees, those built from the shortest possible contacts, effectively freezing the system into a high-affinity backbone. As *kT* rises, the weight distribution flattens, and the ensemble begins to explore an increasingly diverse set of spanning trees. This redistribution is the physical origin of the observed heat capacity peaks and the *kT* -dependent entropy profiles; it confirms that the model captures a genuine thermodynamic ensemble rather than a topologically static description. The cutoff *d*_*c*_ enters only once, to fix which edges exist. Temperature never adds or removes edges; it changes only their relative statistical importance within the fixed topology.

As a concrete illustration, at *kT*_ref_ = 1.0 Å a backbone contact (*d* = 3.8 Å) carries weight *w* ≈ 0.022, whereas a peripheral contact (*d* = 7.9 Å) carries *w* ≈ 3.7 × 10^−4^ roughly two orders of magnitude smaller.

#### Effective temperature

The physical contact energy is *α* · *d*_*ij*_, where *α* converts angstroms to energy. Defining *kT*_eff_ ≡ *k*_*B*_*T/α* absorbs *α* into the temperature scale, giving *kT*_eff_ units of Å. We write *kT* for brevity throughout. Because all state comparisons are formed as differences, they are independent of *α* and carry unambiguous thermodynamic content. Mapping *kT* onto a Kelvin scale would require calibrating *α* against experimental observables (e.g. crystallographic *B*-factors or NMR order parameters), which is not attempted here.

#### Physically sensible window

Two constraints define a reference window within which the Boltzmann weights most closely approximate the contact hierarchy expected for a well-folded protein at ambient conditions. The partition function is, however, analytically exact across the full temperature range; thermodynamic features outside this window reflect genuine properties of the network energy landscape, in particular the competition among near-degenerate low-energy spanning trees that is inaccessible to single-minimum descriptions. To bracket the useful temperature range, we impose two constraints on the Boltzmann weights. Core contacts (*d* ≈ 4 Å) should retain at least 1% of unit weight, ensuring that the shortest interactions remain thermodynamically active; this gives *kT* > 0.87 Å. Peripheral contacts (*d* ≈ 8 Å) should carry less than 1% of unit weight, preventing the ensemble from flattening into the topology-only (GNM) limit; this gives *kT <* 1.74 Å:

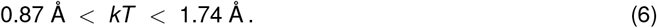

Within this window, short contacts dominate while multiple spanning trees compete, the regime expected for a well-folded protein. The reference value *kT*_ref_ = 1.0 Å sits comfortably inside. The full temperature sweep (*kT* ∈ [0.3, 6.0] Å, 60 uniform grid points) extends well beyond this window in both directions, exploring regimes from single-tree dominance to near-uniform tree weights.

### 2.4 Weighted Kirchhoff Laplacian

The Boltzmann weights are assembled into an *n* × *n* weighted Kirchhoff Laplacian **Γ**(*kT*),

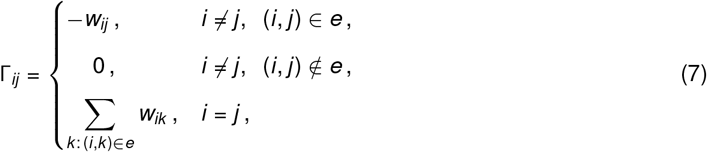

or equivalently **Γ** = **D** − **W**, where **W** is the weighted adjacency matrix and **D** the diagonal matrix of weighted degrees. The matrix is real, symmetric, and positive semidefinite by construction, with a single zero eigenvalue (eigenvector: all-ones) as long as *G* is connected. Previous applications of the graph Laplacian to protein networks have focused on individual eigenvalues as structural descriptors; ^15^ here, the product of all nonzero eigenvalues, the determinant of the reduced Laplacian, gives the partition function exactly.

### 2.5 Spanning-Tree Partition Function

The sum of the edge energies of a spanning tree is:

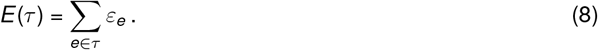

It is this tree-level energy, not any single-edge quantity.

The weighted Matrix-Tree Theorem ^17,18^ provides three equivalent expressions for the partition function:

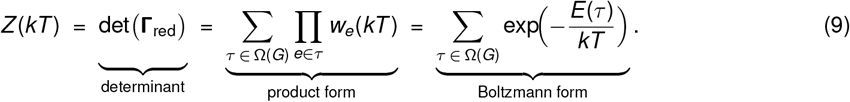

The physical mechanism underlying the temperature-dependent redistribution of statistical weight is illustrated with a minimal four-node graph in Figure S4, where explicit enumeration of spanning trees clarifies the origin of the heat capacity peak and the entropy rise. Here Ω(*G*) is the set of all spanning trees of *G* and **Γ**_red_ is the reduced Laplacian obtained by deleting any one row and the corresponding column, the choice does not affect the result. The first form is what we actually compute: a single determinant replaces the combinatorial sum. The second makes each tree’s contribution explicit as a product of edge weights. The third is the canonical statistical-mechanical form, identifying every spanning tree *τ* as a microstate weighted by its total energy at inverse temperature *β* = 1/(*kT*). It is this identification that turns a graph-theoretic determinant into a bona fide partition function.

Because *G* is connected, **Γ**_red_ is positive definite and *Z* (*kT*) > 0 for all *kT* > 0. For numerical stability, we work with the logarithm,

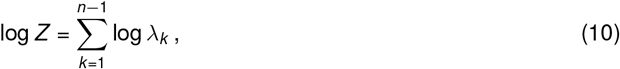

where *λ*_1_, …, *λ*_*n*−1_ are the eigenvalues of **Γ**_red_, computed. Only eigenvalues exceeding 10^−10^ enter the sum, guarding against numerical zeros.

### 2.6 Thermodynamic Quantities

All thermodynamic observables derive from log *Z* (*kT*). Because *Z* (*kT*) is a finite sum of positive exponentials; it is smooth on (0, ∞); all quantities below are therefore real-analytic in *kT* and possesses derivatives of every order.

#### Free energy

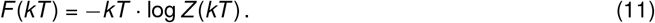

#### Mean tree energy

The Boltzmann-weighted average *Ē* (*kT*) of *E* (*τ*) over all spanning trees,

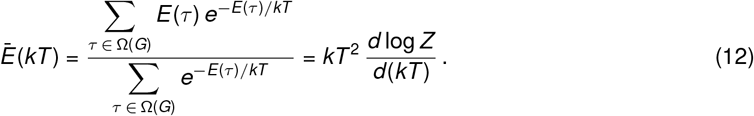

Numerically, we evaluate the right-hand form by a symmetric finite difference with a relative step *δ* = 10^−4^ · *kT* :

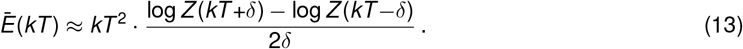

#### Entropy (units of *k*_*B*_)

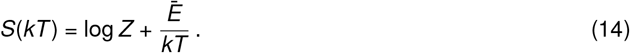

#### Fundamental relation

These quantities satisfy

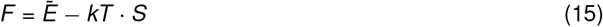

identically an algebraic consequence of Eqs. (11)-(14), not an approximation. Verifying that Eq. (15) holds to machine precision at every grid point is the single most informative test of the computation.

#### Heat capacity

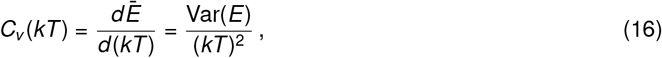

where Var(*E*) is the variance of *E* (*τ*) over the Boltzmann ensemble. The second form is the fluctuation– dissipation relation; because a variance cannot be negative, it guarantees *C*_*v*_ ≥ 0 everywhere.

#### A note on absolute values

Thermodynamic potentials are defined up to an additive constant, so their individual absolute values have no physical meaning. Everything reported in this work is either a state difference Δ*X* = *X*_active_ − *X*_inactive_ or a temperature derivative.

### 2.7 Edge Marginal Inclusion Probabilities

How important is a given contact for holding the network together? The marginal probability that edge *e* = (*i, j*) appears in a randomly sampled spanning tree answers this question:

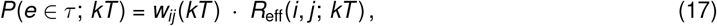

Where

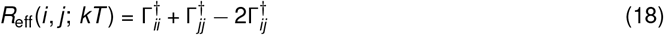

is the effective resistance between the two nodes in the conductance network {*w*_*e*_}_*e*∈*E*_, and **Γ**^†^ is the Moore-Penrose pseudoinverse of **Γ**. ^20,21^ In the GNM, the pseudoinverse element (**Γ**^−1^)_*ij*_ equals the mean-square fluctuation 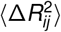 between residues *i* and *j*, a quantity termed the dynamic distance. ^13^ The effective resistance *R*_eff_(*i, j*) used here (Eq. 18) generalises this relationship to the weighted Laplacian: in the unweighted limit (*kT* → ∞). The two quantities coincide, establishing a direct correspondence between the network-theoretic measure of signal propagation and the GNM description of correlated residue motion.

In practice we compute **Γ**^†^ by inverting the reduced Laplacian **Γ**_red_ (with `scipy.linalg.inv`) and embedding the result into an *n* × *n* matrix whose ground-node row and the column is zero. All marginal probabilities are clipped to [0, 1] afterwards to absorb floating-point noise.

A high *P*(*e* ∈ *τ*) means the contact is structurally essential; it shows up in most spanning trees, not merely that the two residues happen to be close. The identity

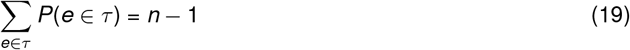

holds by construction (every tree has *n* − 1 edges, so the expected count must be *n* − 1) and serves as a second independent check on the numerics. Inclusion probabilities are reported at *kT*_ref_ = 1.0 Å.

### 2.8 Relationship to the Gaussian Network Model

The present framework and the standard GNM share the same underlying graph *G*: both connect residues within a cutoff *d*_*c*_ and ignore everything beyond it. The difference lies entirely in the edge weights. The GNM sets *w*_*ij*_ = 1 for every contact; we set *w*_*ij*_ = exp(−*ε*_*ij*_ */kT*).

The GNM is therefore a limiting case. As *kT* → ∞ every Boltzmann weight approaches unity,

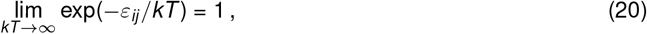

and the weighted Laplacian converges to the unweighted one: **Γ**(*kT*) → **Γ**^GNM^. The partition function becomes the spanning-tree count, *Z* → |Ω(*G*)| is the number of spanning trees, and the thermodynamic quantities collapse according

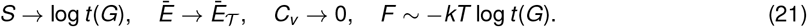

where 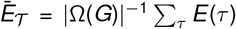 is the unweighted mean tree energy. The entropy saturates at a purely topological invariant; the heat capacity vanishes because all trees are now equiprobable and there is nothing left to discriminate among them. Similarly, the edge inclusion probabilities reduce to the unweighted effective resistances: 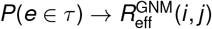.

This limit is more than a theoretical curiosity; it provides an internal consistency check. At high enough *kT*, every quantity computed from the weighted ensemble must converge to its GNM counterpart; failure to do so would indicate a bug.

### 2.9 Functional Region Classification

Residues are grouped into functional regions following standard KRAS annotations (Table 1). Inter-lobe bridge contacts are edges (*i, j*) with one endpoint in the effector lobe (*i* ≤ 86) and the other in the allosteric lobe (*j* ≥ 87).

**Table 1:** KRAS functional region definitions.

| Region | Residues |
| --- | --- |
| P-loop | 10-17 |
| Switch I | 25-40 |
| Switch II | 57-75 |
| Effector lobe (other) | 1-86 (excl. above) |
| Allosteric lobe | 87-169 |
| C-terminal | 170-172 |

**Table 2:**
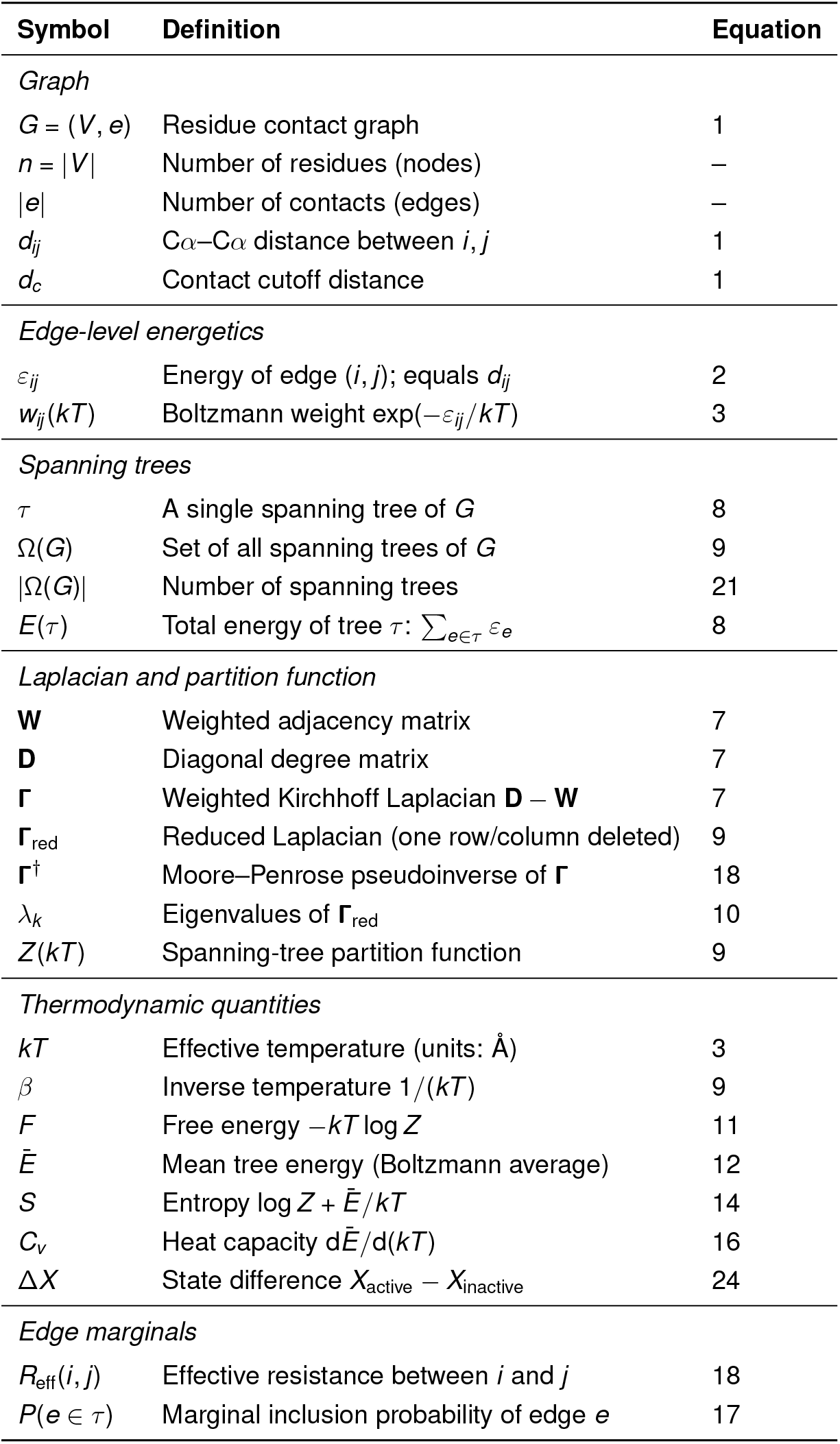
Symbol definitions used throughout this work.

**Table 3:**
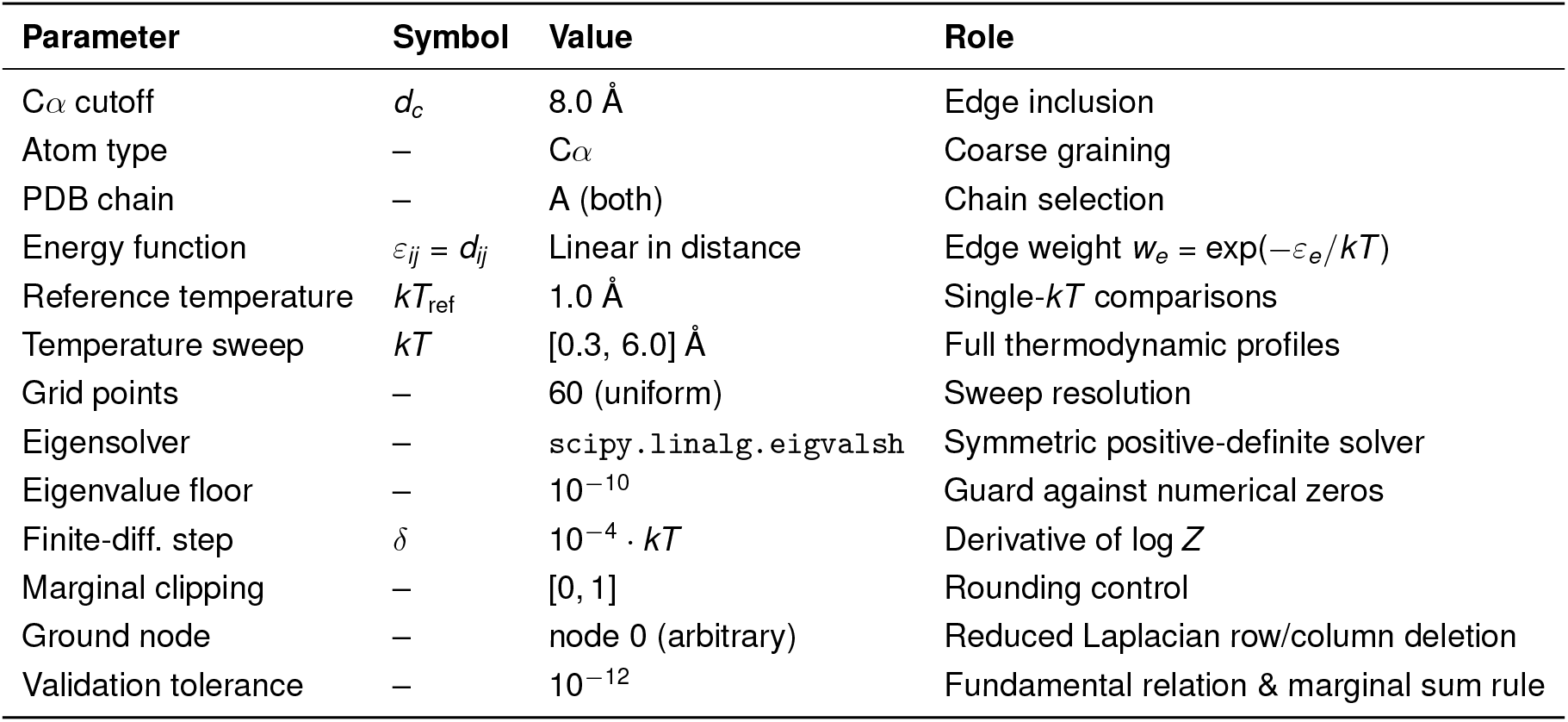
Complete parameter specification.

| Parameter | Symbol | Value | Role |
| --- | --- | --- | --- |
| Cα cutoff | $d_c$ | 8.0 Å | Edge inclusion |
| Atom type | – | Cα | Coarse graining |
| PDB chain | – | A (both) | Chain selection |
| Energy function | $\varepsilon_{ij} = d_{ij}$ | Linear in distance | Edge weight $w_e = \exp(-\varepsilon_e/kT)$ |
| Reference temperature | $kT_{\text{ref}}$ | 1.0 Å | Single-$kT$ comparisons |
| Temperature sweep | $kT$ | [0.3, 6.0] Å | Full thermodynamic profiles |
| Grid points | – | 60 (uniform) | Sweep resolution |
| Eigensolver | – | `scipy.linalg.eigvalsh` | Symmetric positive-definite solver |
| Eigenvalue floor | – | $10^{-10}$ | Guard against numerical zeros |
| Finite-diff. step | $\delta$ | $10^{-4} \cdot kT$ | Derivative of $\log Z$ |
| Marginal clipping | – | [0, 1] | Rounding control |
| Ground node | – | node 0 (arbitrary) | Reduced Laplacian row/column deletion |
| Validation tolerance | – | $10^{-12}$ | Fundamental relation & marginal sum rule |

### 2.10 Numerical Validation

Four checks guard against implementation errors. The absolute thermodynamic quantities for the active and inactive states are shown in Figure S1; they satisfy the relation *F* = *Ē* − *TS* to numerical precision throughout the entire temperature range.

1. **Fundamental relation**. |*F* − (*Ē* − *kT S*)| *<* 10^−12^ at every grid point (Eq. 15).
2. **Marginal sum rule**. 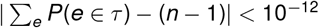 (Eq. 19).
3. **Non-negative heat capacity**. *C*_*v*_ ≥ 0 everywhere, as guaranteed by the fluctuation–dissipation form (Eq. 16).
4. **Moore–Penrose throughout**. The pseudoinverse **Γ**^†^ satisfies

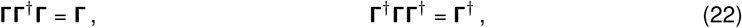

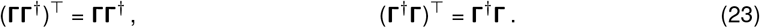

All four passed with residuals below 10^−12^ for both structures at every temperature.

### 2.11 Notation

### 2.12 Parameters

### 2.13 Implementation

Everything was coded in Python 3 and run inside Google Colaboratory (2-core CPU, 12 GB RAM). The libraries involved are NumPy (array operations), SciPy (eigendecomposition via `scipy.linalg.eigvalsh;` inversion via `scipy.linalg.inv`), NetworkX (graph construction and traversal), and Matplotlib (figures). The full pipeline, PDB parsing through final figures, is deterministic and finishes in under two minutes. No external simulation software is needed.

Contact maps (Figures 3, 4) use `imshow` with the `YlOrRd` colormap for absolute probabilities and `RdBu_r` for differential maps; colour limits are clipped to the 95th percentile of absolute values. Thermodynamic profiles (Figures 1, 2) overlay shaded spans for the functional regions.

**Figure 1:**
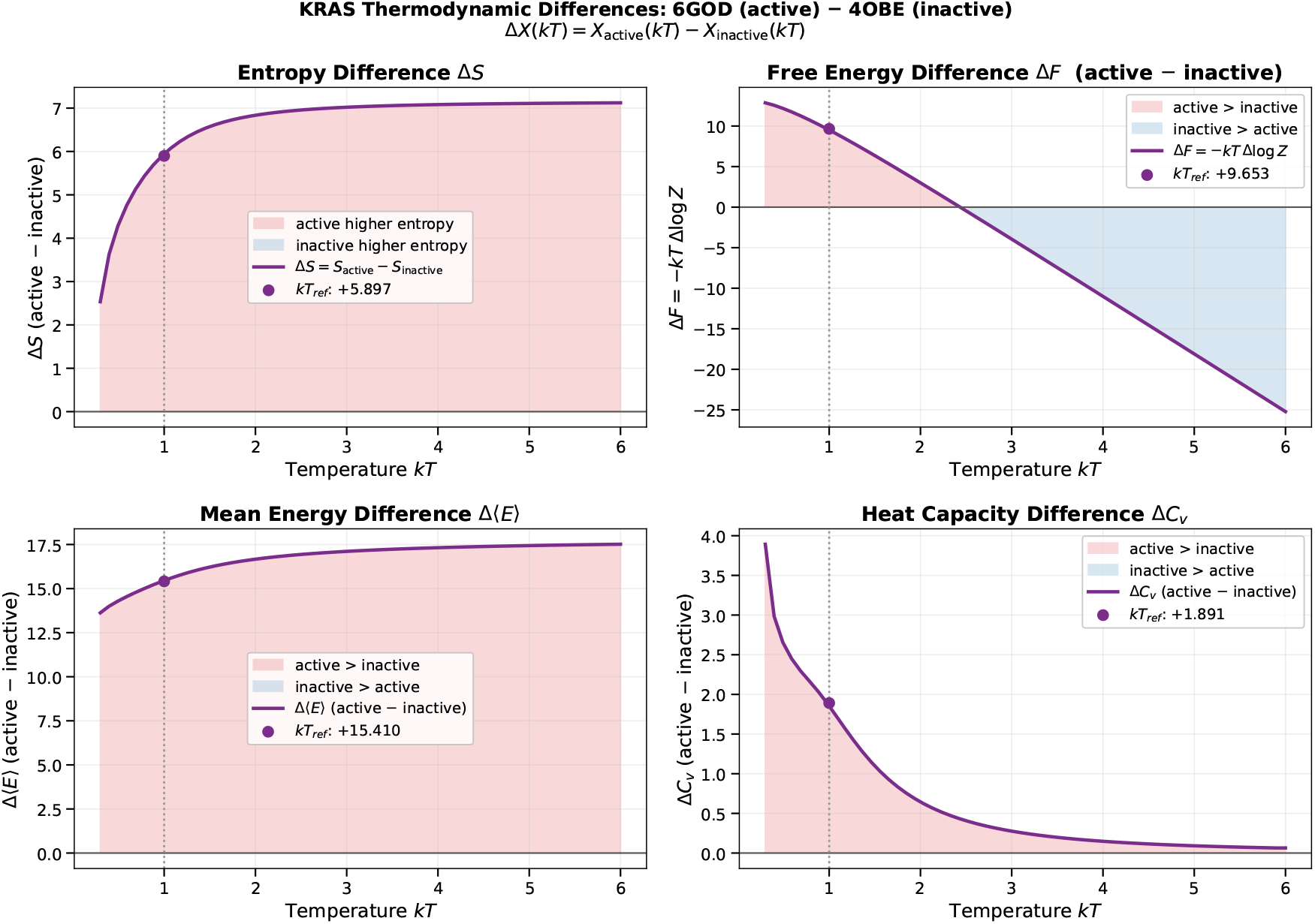
Thermodynamic difference profiles. (*kT*_ref_ = 1.0, dotted line). Δ*X* (*kT*) = *X*_active_(*kT*) − *X*_inactive_(*kT*) for all four observables across *kT* ∈ [0.3, 6.0]. **(A)** Entropy difference Δ*S*: uniformly positive and saturating near Δ*S*≈ 7 at high temperature, confirming that the GTP-bound network supports more spanning-tree configurations at every *kT* . At *kT*_ref_ = 1.0 Å, Δ*S* = +5.76. **(B)** Free energy difference Δ*F* = − *kT* Δlog *Z* : positive at physiological temperatures, crossing zero near *kT*≈ 2.4. **(C)** Mean tree energy difference Δ*Ē* > 0 throughout: the active network pays a persistent energetic cost reflecting the structural reorganisation required for effector engagement. **(D)** Heat capacity difference Δ*C*_*v*_ : large at low temperature and decaying monotonically, indicating greater thermal sensitivity of the active-state network.

**Figure 2:**
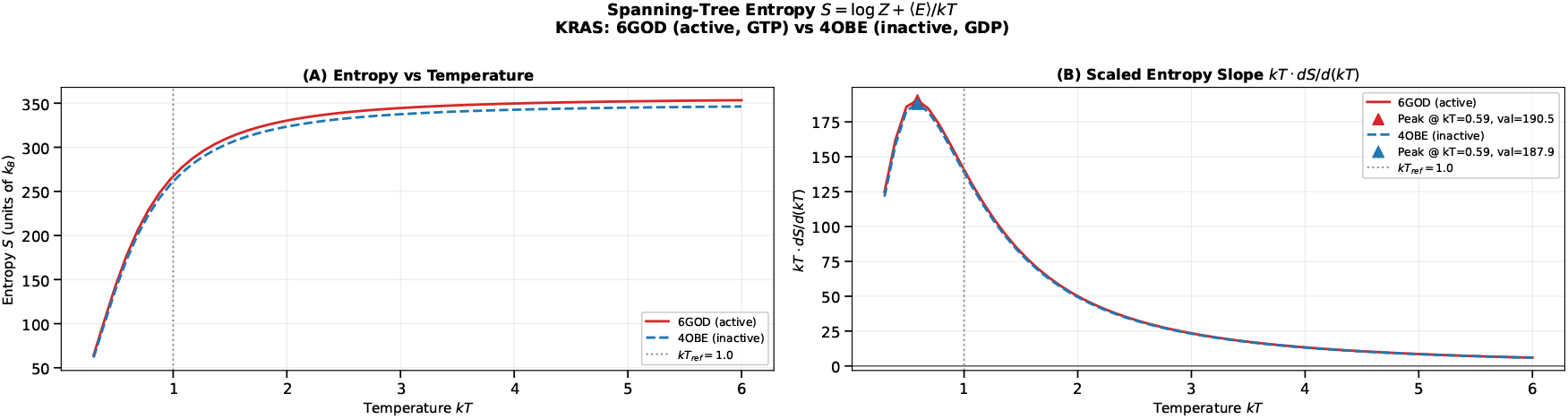
Temperature dependence of spanning-tree entropy. **(A)** Absolute entropy (14) (units of *k*_*B*_) for 6GOD (red solid) and 4OBE (blue dashed). Both curves increase monotonically and saturate at high temperature, with the active state consistently above the inactive state, indicating higher configurational degeneracy of the GTP-bound contact network. **(B)** Scaled entropy slope *kT*· *dS/d* (*kT*). Both states peak at *kT*≈ 0.59 Å (6GOD: 190.5 *k*_*B*_; 4OBE: 187.9 *k*_*B*_), showing that the rate of topological reorganisation is maximal, well below *kT*_ref_ = 1.0 Å, and that the two networks reorganise at nearly identical rates despite their different nucleotide states. The dotted vertical line marks *kT*_ref_.

**Figure 3:**
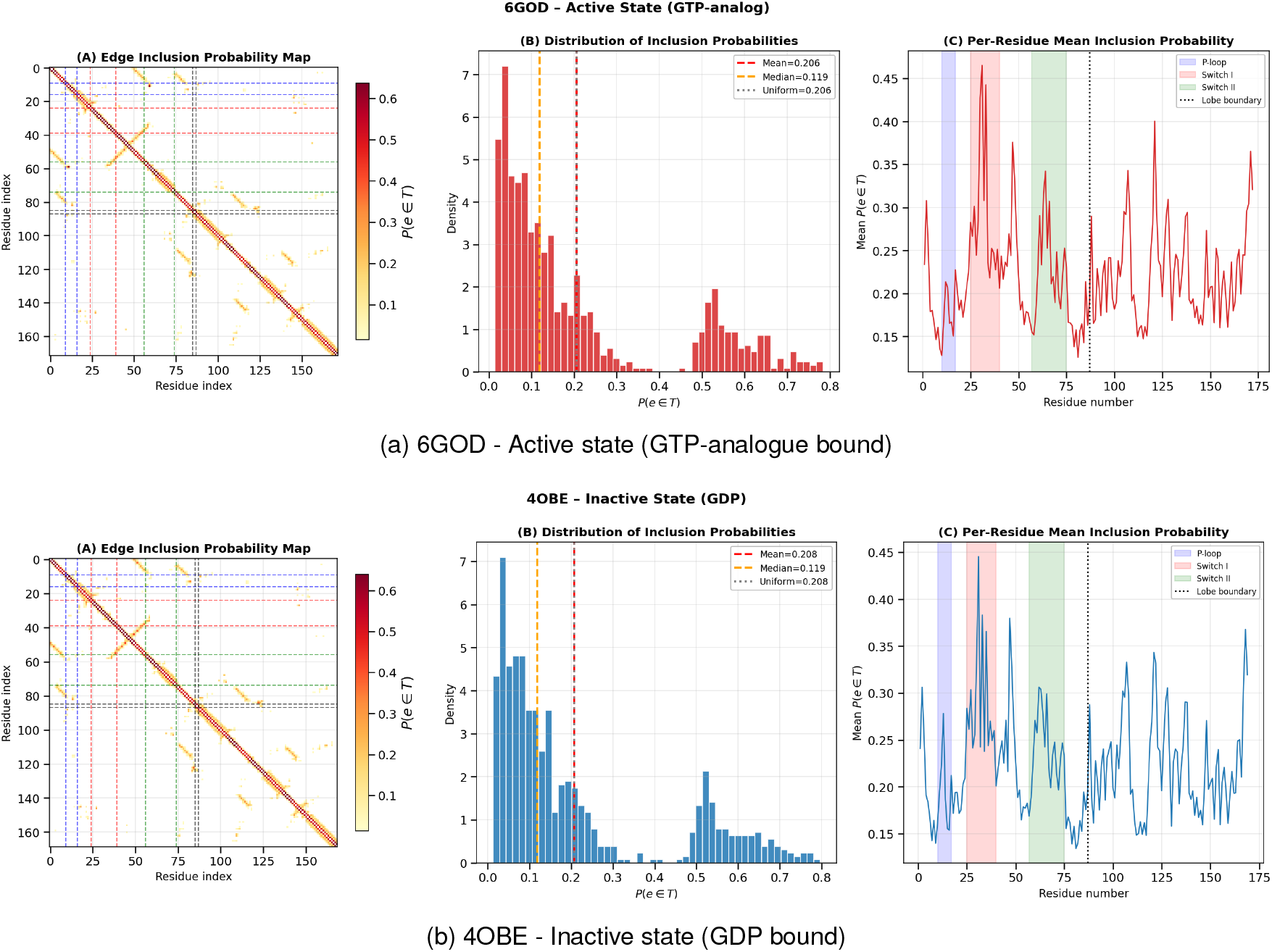
Edge marginal inclusion probability maps. Three-panel layout for each state.**(A)** Contact map coloured by *P* (*e* ∈ *T*) (yellow-red scale, vmin = 0, vmax = 1). **(B)** Histogram of inclusion probabilities with mean (red dashed), median (orange dashed), and uniform reference (gray dotted). **(C)** Per-residue mean *P*(*e* ∈ *T*) with functional regions shaded: P-loop (blue, residues 10-17), Switch I (red, 25-40), Switch II (green, 57-75), and lobe boundary (dotted, residue 87).

**Figure 4:**
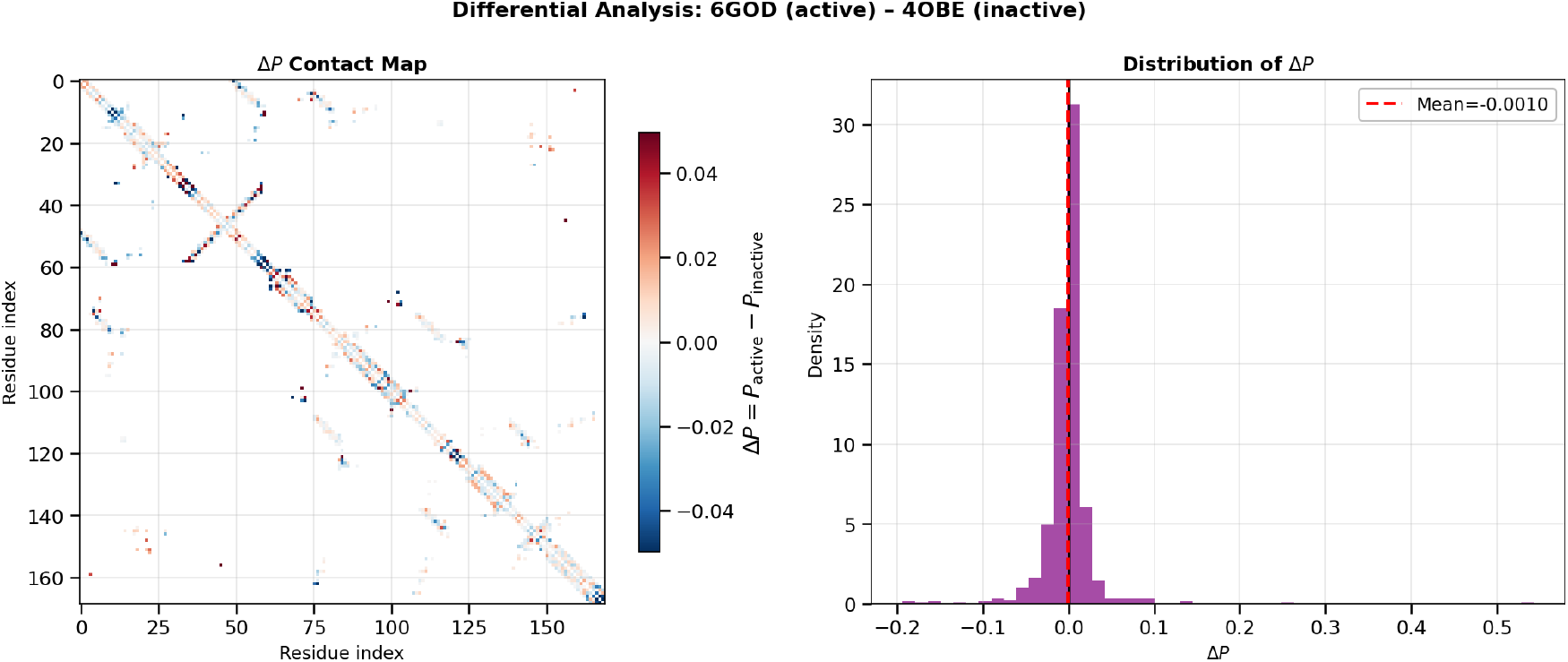
Differential edge inclusion probability analysis. **(Left)** Δ*P* contact map (diverging red-blue colour scale; red = active enriched, blue = inactive enriched). Differences are sparse and localised, concentrated in Switch I and at the effector, allosteric lobe interface. **(Right)** Distribution of Δ*P* values across 783 common edges; mean = −0.001, confirming marginal conservation of total spanning-tree weight between states.

## 3. RESULTS

### 3.1 Network Statistics

Table 4 summarises the basic topological properties of the two contact networks. The active structure 6GOD contains three additional residues and 21 additional contacts relative to 4OBE, reflecting the difference in resolved residues between the two crystals. Despite this size difference, mean and median edge marginal probabilities *P*(*e* ∈ *τ*) are nearly identical (mean 0.206 versus 0.208; median 0.119 in both cases), indicating that the average structural essentiality of a contact is conserved between states. This near-invariance implies that the two networks are comparably well-connected and that thermodynamic differences between them arise not from changes in overall connectivity density but from the redistribution of structurally critical contacts across the network.

**Table 4:** Contact network statistics for the two KRAS structures at C_*α*_ cutoff 8.0 Å.

| Property | 6GOD (active, GTP) | 4OBE (inactive, GDP) |
| --- | --- | --- |
| Nucleotide state | GTP-analogue bound | GDP bound |
| Residues (nodes) | 172 | 169 |
| Contacts (edges) | 830 | 809 |
| Mean $P(e \in T)$ | 0.206 | 0.208 |
| Median $P(e \in T)$ | 0.119 | 0.119 |

### 3.2 Thermodynamic Difference Profiles

Because individual thermodynamic potentials are defined only up to an additive constant (§ 2.6), the physical content resides entirely in the state differences

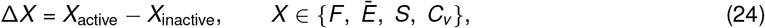

which are shown in Figure 1. The free-energy difference Δ*F* crosses zero at *kT* ≈ 2.41 Å, a crossover independent of any additive reference constant, marking the effective temperature at which the active and inactive ensembles are thermodynamically balanced. A complete panel of individual quantities, their differences, and the heat capacity is provided in Figure S2, offering a detailed view of the thermodynamic behavior of both states. Below this value, the inactive state is preferred; above it, the active state becomes favoured. This balance, and its perturbation by nucleotide binding, underlies the switching mechanism.

### 3.3 Temperature Dependence of Entropy and Heat Capacity

That the heat-capacity maximum lies below the reference window is structurally expected. The peak reflects the smallest energy scale in the contact network: the energetic cost of substituting one or two edges in the ground-state spanning tree with slightly longer alternatives, a local intra-state property of the protein’s core packing. This scale is distinct from the interstate switching scale captured by Δ*F*, Δ*S*, and Δ*Ē*, which operates within and above the reference window.

The framework therefore resolves two thermodynamic regimes simultaneously: core contact competition at low *kT* and global nucleotide-state discrimination at moderate *kT* .

Figure 2 shows the thermodynamic entropy

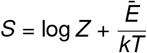

for both states as a function of effective temperature. The active state maintains higher entropy across the entire range (Δ*S* > 0), saturating near Δ*S* ≈ 7 at high *kT* . This persistent advantage reflects the greater topological degeneracy, i.e., the larger number of viable spanning trees of the GTP-bound network. In the high-temperature limit, the entropy of each state converges to log *t* (*G*) (Eq. 21), confirming that the saturation value is a topological invariant. The scaled entropy slope *kT* d*S*/d(*kT*) (Figure 2B) peaks sharply at low temperature and converges for both states at high temperature, indicating that thermal reorganisation of the two networks becomes indistinguishable above *kT* ≈ 3 Å.

The heat capacity

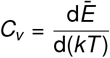

reaches a pronounced maximum near *kT* ≈ 0.53 Å, below the physically sensible window of Eq. (6) (0.87 Å < *kT* < 1.74 Å). This feature reflects the activation of the first excited spanning trees above the network’s ground state.

At very low temperatures, the Boltzmann weights *w*_*ij*_ = exp(−*d*_*ij*_ /*kT*) are overwhelmingly dominated by the shortest edges, and the ensemble collapses onto a single dominant tree, the unique connected subgraph composed of the shortest possible edges reaching every residue. As the temperature rises, a small number of slightly longer edges acquire sufficient weight to replace one or two edges in the ground-state tree, generating a handful of low-lying excited configurations. The heat capacity peak marks the point of maximal competition between the ground state and these first alternatives. That *C*_*v*_ → 0 at high *kT* (Eq. 21) confirms that the peak is a finite-temperature phenomenon: once all trees are equiprobable, the variance of the tree energy vanishes, and the heat capacity disappears.

That this peak lies below the physically sensible window is natural. It reports on the smallest length scale in the contact network: the difference between the very shortest edges and those that can replace them, typically on the order of 1-2 Å. The heat capacity maximum, therefore, reflects local, high-fidelity core contacts defining the protein’s ground-state architecture, whereas global switching between active and inactive states involves much larger effective energy scales supplied not by temperature but by nucleotide binding.

### 3.4 Edge Inclusion Probabilities

Marginal edge inclusion probabilities *P*(*e* ∈ *T*) at *kT*_ref_ = 1.0 Å are shown in Figure 3 for both states. The distributions are broad and right-skewed, peaking near *P* ≈ 0.05 with a secondary shoulder near *P* ≈ 0.5-0.7. Most contacts are topologically redundant; a minority serve as essential backbone edges. The mean *P* equals the uniform reference (*n* − 1)*/*|*E* |, confirming consistency with the sum rule (Eq. 19).

This localisation is structurally expected. Switch I (residues 25-40) constitutes the direct binding interface for downstream effectors including Raf-RBD and PI3K*γ*, and its conformational state gates signal transduction to the MAPK and PI3K cascades. That Switch I emerges as the dominant locus of state-dependent Δ*P* provides an internal validation of the method, recovering the established primary site of nucleotide-driven functional reorganisation from network topology alone.

### 3.5 Differential Contact Analysis

Figure 4 presents the differential map Δ*P* = *P*_active_ − *P*_inactive_ for the 783 contacts common to both structures. The mean Δ*P* = −0.001 is near zero by construction; the informative signal lies in the spatial distribution of deviations. Switch I (residues 25-40) dominates the positive Δ*P* values (contacts more essential in the active state), followed by the inter-lobe region near residues 86-90. Negative Δ*P* contacts, those more essential in the inactive state, concentrate at the Switch II/allosteric lobe interface.

## 4 PRINCIPAL FINDINGS

The following analysis is based on a single representative crystal structure per nucleotide state. It should therefore be interpreted as a proof-of-concept demonstration of the framework rather than a definitive characterization of the KRAS thermodynamic landscape.

The thermodynamic distinction between the active and inactive states of KRAS can be distilled into three quantities: the free energy difference Δ*F*, the mean tree energy difference Δ*Ē*, and the conformational entropy difference Δ*S*. Taken together, they reveal how the protein’s contact network is rewired upon nucleotide exchange, and why this rewiring matters.

### Free energy and the thermodynamic ground state

At physiological temperatures, the inactive state is thermodynamically favoured, carrying lower free energy than the active state (Δ*F* > 0). The GDP-bound conformation is structurally well integrated: Switch I and Switch II lie collapsed against the protein core, forming a dense, stable network of contacts. Without an external driving force, the system therefore predominantly occupies the inactive basin. GTP binding supplies the free energy required to populate the higher-free-energy active conformation.

The *intrinsic* balance point, where neither state is thermodynamically preferred, occurs at *kT* ≈ 2.41. Unlike the zero-crossings of the individual free energy curves, which both fall near *kT* ≈ 2.2 as an artefact of the unit convention (see § 2.6), this crossover is independent of any additive reference. That it lies well above the physiological effective temperature underscores the role of the inactive state as the intrinsic ground state and confirms that activation requires sustained nucleotide-derived free energy.

### Energetic cost of activation

Decomposing the free energy difference clarifies its origin. The active state exhibits a consistently higher mean tree energy (Δ*E* > 0): its residue-residue contacts are, on average, less optimal than those in the inactive state. In the GDP-bound structure, the switches are held in place by favourable interactions with the core and with each other; GTP binding disrupts many of these, forcing the switches into extended and partially exposed conformations. The resulting energetic cost is not a defect but a necessary investment, the price of creating an accessible effector-binding surface.

### Entropic gain and functional versatility

What justifies this investment is the accompanying gain in conformational entropy. The active state has higher entropy (Δ*S* > 0), indicating that its switch regions are not simply rearranged but access a larger number of competing network configurations. In the inactive state, the switches are confined to a narrow ensemble of low-energy spanning trees; GTP binding liberates them, granting access to a much broader set of thermally accessible configurations. In network terms, this entropic expansion corresponds to a larger number of topologically distinct spanning trees connecting the nucleotide-binding pocket to distal surface residues. The formalism establishes this combinatorial diversity directly; its biological correlate, that a richer set of network configurations facilitates engagement of multiple downstream effectors including RAF, PI3K, and RALGDS through distinct allosteric routes, remains an interpretive hypothesis consistent with the present analysis but requiring validation against conformational and binding data.

### Temperature dependence and the switching mechanism

The temperature dependence reinforces this picture. As temperature rises, the entropic term *T* Δ*S* grows, gradually offsetting the energetic penalty Δ*Ē* . At sufficiently high effective temperatures, the two states become equally probable (Δ*F* = 0). That the crossover lies well above physiological temperatures reaffirms that the inactive state is the intrinsic ground state and that switching demands sustained nucleotide-derived free energy.

KRAS activation is therefore not a simple flip from a low-energy off state to a high-energy on state. It is a redistribution of thermodynamic resources: the energetic cost of disrupting ground-state contacts is offset by a gain in conformational entropy, which in turn enables functional versatility. The active state is structurally diverse, and it is this diversity encoded in the network architecture that may enable a single protein to serve as a hub for multiple signalling pathways.

## 5 DISCUSSION

MD simulations by Zhao et al. ^6^ show that GTP binding enhances conformational flexibility and long-range interactions in KRAS, consistent with the higher entropy we observe in the active-state network. Their analysis centres on pairwise correlations and kinetic pathways; our framework quantifies the global network consequences, showing that the entropic gain comes at a persistent energetic cost Δ⟨*E* ⟩ > 0 and that the two states are tuned to operate near a thermodynamic crossover that maximises switch sensitivity.

Allosteric network modelling of KRAS–effector interactions using Markov state models and mutational scanning reveals that oncogenic mutations redistribute communication pathways between the nucleotide-binding pocket and the effector interface. ^9^ The large Δ*P* we observe in Switch I accords with this picture and provides a direct thermodynamic measure of the redistribution. That a small number of contacts can dominate this process is expected from the small-world topology of protein contact networks, ^14^, which ensures that a few edges serve as critical bridges for long-range signal transmission. Edges with high *P*(*e* ∈ *τ*) in our analysis are precisely these bridges: they appear in the majority of spanning trees and constitute the most robust communication channels.

Crystallographic work by Shima et al. ^24^ established that the GTP-bound form of Ras exists as an ensemble of two interconverting conformations (states 1 and 2), with the switch regions exhibiting pronounced flexibility, suggesting that conformational entropy plays a key role in activation. Our framework quantifies this entropic contribution directly from network topology.

Pálfy et al. ^25^ highlighted the critical role of the Mg^2+^-free state in facilitating nucleotide exchange in oncogenic KRAS mutants, underscoring the importance of conformational flexibility in the switch regions. Our analysis complements their findings: the active state’s higher entropy (Δ*S* = +5.76) reflects the increased conformational freedom of the switches upon GTP binding, a freedom likely coupled to Mg^2+^ release during nucleotide exchange. The underlying molecular mechanism of GTP hydrolysis has been characterised in atomic detail by time-resolved FTIR spectroscopy. ^26^

MD studies by Chen et al. ^27^ align with our finding of higher tree energy in the active state, directly showing that key contacts—particularly between GTP and Switch I weaken or become unstable upon activation. Xu et al. ^28^ provide complementary structural evidence: although a new interaction forms (Tyr32 with the GTP *γ*-phosphate), stable pre-existing contacts are disrupted. The overall picture supports our conclusion that the active state is characterised by less stable packing and a higher-energy contact network compared to the tightly coordinated inactive conformation.

Senet et al. ^19^ showed that Shannon entropy computed over protein subgraphs at graph distance *D* = 1 exhibits a sharp transition near the melting temperature in gpW and varies heterogeneously along the sequence of *α*-synuclein. Their local entropy captures the diversity of micro-environments around each residue; ours captures the combinatorial degeneracy of the entire network. Both derive from the Kirchhoff Laplacian, and both demonstrate that graph-based entropy can discriminate functionally distinct protein states, at the residue level in their case, at the network level in ours.

The entropic basis of activation bears directly on the inhibitor mechanism. Covalent inhibitors such as sotorasib and adagrasib function by trapping KRAS in the GDP-bound inactive conformation; within the present framework, this admits a network-level interpretation: these compounds effectively collapse the spanning-tree ensemble toward the low-entropy inactive state, suppressing the combinatorial degeneracy (Δ*S* > 0) that enables engagement of multiple downstream effectors. Whether spanning-tree multiplicity constitutes a useful metric for prospective inhibitor design remains to be tested against binding data and mutant-specific structural ensembles.

A limitation of this analysis is its reliance on two static crystallographic structures, which do not capture the full conformational heterogeneity accessible to each nucleotide state. Effects of crystal packing, incomplete loop resolution, and the absence of explicit solvent are not represented. Extending the framework to conformational ensembles derived from molecular dynamics trajectories, or to multiple independently determined crystal structures per state, would provide a more stringent test of these conclusions and constitutes a natural direction for future work.

## 6 CONCLUSION

We have applied a spanning-tree partition function analysis to two KRAS crystal structures, revealing thermodynamic differences that are coherent and biologically interpretable. The active (GTP-bound) state carries a higher energetic cost but a larger conformational entropy, with the intrinsic free-energy balance crossing at *kT* ≈ 2.41 Å; Switch I emerges as the primary locus of state-dependent topological change. These findings support a view of KRAS activation as an entropy-driven investment: GTP binding pays an energetic price to liberate the switch regions into a diverse ensemble of competing networks configurations.

For KRAS specifically, natural extensions include application to oncogenic mutants (G12C, G12D, G12V), where mutation-specific shifts in the entropy and enthalpy balance may provide a thermodynamic rationale for differential drug sensitivity, and calibration of the effective temperature scale against molecular dynamics ensembles or experimentally determined *B*-factors.

More broadly, the framework requires nothing beyond atomic coordinates and a contact cutoff; it yields exact thermodynamic potentials from a single matrix determinant without sampling, force fields, or normal-mode approximations. Any protein for which a residue contact graph can be constructed, including multi-domain signalling hubs, allosteric enzymes, and intrinsically disordered regions upon complex formation, is accessible to the same analysis. The KRAS application presented here demonstrates the approach; the method itself is general.

## AUTHOR CONTRIBUTIONS

**Fatma Senguler Ciftci:** Software, Formal analysis, Investigation, Data curation, Visualization, Writing, original draft. **Burak Erman:** Conceptualization, Methodology, Supervision, Writing, review & editing.

## DATA AVAILABILITY STATEMENT

All source code used for constructing the residue-contact graphs, evaluating the spanning-tree partition functions, and computing the thermodynamic quantities (*F, Ē, C*_*v*_, and *S*) is publicly available on GitHub at https://github.com/fatmasenguler/KRAS_spanning_tree. The repository contains the primary Python scripts, Jupyter notebooks, and a detailed `README` for reproducing the results. The crystallographic structures analyzed in this study (PDB IDs: 6GOD and 4OBE) can be obtained from the RCSB Protein Data Bank (https://www.rcsb.org).

## ACKNOWLEDGEMENTS

Gemini 3 Flash was used during development to assist with Python linear algebra routines (Laplacian construction, pseudoinversion) and to polish English grammar. All code and text were reviewed, edited, and validated by the authors, who bear full responsibility for the final manuscript.

## SUPPORTING INFORMATION

Additional supporting information may be found online in the Supporting Information section at the end of this article.

**Figure S1**. Individual thermodynamic quantities for active (6GOD) and inactive (4OBE) KRAS networks.

**Figure S2**. Comprehensive nine-panel comparison of KRAS thermodynamic states.

**Figure S3**. Sensitivity of the meaningful *kT* range to the contact-distance cutoff.

**Figure S4**. Spanning-tree thermodynamics illustrated on a four-node toy graph.

## References

[1] Prior IA, Hood FE, Hartley JL. The frequency of Ras mutations in cancer. Cancer Res. 2012;72:2457–2467.

[2] Grant BJ, Gorfe AA, McCammon JA. Ras conformational switching: simulating nucleotide-dependent conformational transitions with accelerated molecular dynamics. PLoS Comput Biol. 2009;5:e1000325.

[3] Ostrem JM, Shokat KM. Direct small-molecule inhibitors of KRAS: from structural insights to mechanism-based design. Nat Rev Drug Discov. 2016;15:771–785.

[4] Manley LJ, Lin MM. Kinetic and thermodynamic allostery in the Ras protein family. Biophys J. 2023;122:3882–3893.

[5] Killoran RC, Smith MJ. Conformational resolution of nucleotide cycling and effector interactions for multiple small GTPases determined in parallel. J Biol Chem. 2019;294:9937–9948.

[6] Zhao DR, Yang JT, Liu MT, Yang LQ, Sang P. Deciphering allosteric mechanisms in KRAS activation: insights from GTP-induced conformational dynamics and interaction network reorganization. RSC Adv. 2025;15:2261–2274.

[7] Rennella E, Henry C, Dickson CJ, Georgescauld F, Wales TE, Erdmann D, et al. Dynamic conformational equilibria in the active states of KRAS and NRAS. RSC Chem Biol. 2025;6:106–118.

[8] Alshahrani M, Parikh V, Foley B, Hu G, Verkhivker G. Probing binding and allosteric mechanisms of the KRAS interactions with monobodies and affimer proteins. Phys Chem Chem Phys. 2025;27:11242–11263.

[9] Xiao S, Alshahrani M, Hu G, Tao P, Verkhivker G. Exploring binding and allosteric energy landscapes for the KRAS interactions with effector proteins using Markov state modeling of conformational ensembles and allosteric network modeling. Protein Sci. 2025;34(8):e70228.

[10] Yang L, Song G, Jernigan RL. Protein elastic network models and the ranges of cooperativity. Proc Natl Acad Sci USA. 2009;106:12347–12352.

[11] Park JK, Jernigan RL, Wu Z. Coarse-grained normal mode analysis vs. refined Gaussian network model for protein residue-level structural fluctuations. Bull Math Biol. 2013;75:124–160.

[12] Togashi Y, Flechsig H. Coarse-grained protein dynamics studies using elastic network models. Int J Mol Sci. 2018;19:3899.

[13] Erman B. The Gaussian network model as a framework for allosteric analysis: dynamic distance, edge centrality, and entropy sensitivity in KRAS. Preprint. Posted October 4, 2025. bioRxiv. doi:10.1101/2025.05.18.654696.

[14] Atilgan AR, Akan P, Baysal C. Small-world communication of residues and significance for protein dynamics. Biophys J. 2004;86:85–91.

[15] Gadiyaram V, Vishveshwara S, Vishveshwara S. From quantum chemistry to networks in biology: a graph spectral approach to protein structure analyses. J Chem Inf Model. 2019;59:1715–1727.

[16] Liang Z, Verkhivker GM, Hu G. Integration of network models and evolutionary analysis into highthroughput modeling of protein dynamics and allosteric regulation: theory, tools and applications. Brief Bioinform. 2020;21:815–835.

[17] Kirchhoff G. Über die Auflösung der Gleichungen, auf welche man bei der Untersuchung der linearen Vertheilung galvanischer Ströme geführt wird. Ann Phys. 1847;148:497–508.

[18] Chung FRK. Spectral Graph Theory. Providence, RI: American Mathematical Society; 1997.

[19] Tyler S, Laforge C, Guzzo A, Nicolaï A, Maisuradze GG, Senet P. Einstein model of a graph to characterize protein folded/unfolded states. Molecules. 2023;28:6659.

[20] Lyons R. Asymptotic enumeration of spanning trees. Comb Probab Comput. 2005;14:491–522.

[21] Doyle PG, Snell JL. Random Walks and Electric Networks. Washington, DC: Mathematical Association of America; 1984.

[22] P. J. Flory and B. Erman, “Theory of elasticity of polymer networks. III,” Macromolecules, vol. 15, no. 3, pp. 800–806, 1982.

[23] Bahar I, Atilgan AR, Erman B. Direct evaluation of thermal fluctuations in proteins using a single-parameter harmonic potential. Fold Des. 1997;2:173–181.

[24] Shima F, Ijiri Y, Muraoka S, Liao J, Ye M, Araki M, et al. Structural basis for conformational dynamics of GTP-bound Ras protein. J Biol Chem. 2010;285:22696–22705.

[25] Pálfy G, Menyhárd DK, Ákontz-Kiss H, Vida I, Batta G, Tőke O, Perczel A. The importance of Mg^2+^-free state in nucleotide exchange of oncogenic K-Ras mutants. Chem Eur J. 2022;28:e202201449.

[26] Kötting C, Kallenbach A, Suveyzdis Y, Wittinghofer A, Gerwert K. The GAP-catalyzed GTP hydrolysis reaction of Ras as revealed by time-resolved FTIR spectroscopy. Proc Natl Acad Sci USA. 2008;105:6260–6265.

[27] Chen J, Zhang S, Zeng Q, Wang W, Zhang Q, Liu X. Free energy profiles relating with conformational transition of the switch domains induced by G12 mutations in GTP-bound KRAS. Front Mol Biosci. 2022;9:912518.

[28] Xu S, Long BN, Boris GH, Chen A, Ni S, Kennedy MA. Structural insight into the rearrangement of the switch I region in GTP-bound G12A K-Ras. Acta Crystallogr D Struct Biol. 2017;73:970–984.

[29] Vishveshwara S, Brinda KV, Kannan N. Protein structure: insights from graph theory. J Theor Comput Chem. 2002;1:187–211.

[30] Bagler G, Sinha S. Assortative mixing in protein contact networks and protein folding kinetics. Bioinformatics. 2007;23:1760–1767.

