## Supplementary Information for "Generalizing the Gaussian Network Model: Spanning-Tree Thermodynamics Shows Entropy-Driven KRAS Activation"

This Supporting Information accompanies the main manuscript and contains three supplementary figures referenced therein. All reference numbers correspond to the bibliography of the main article unless otherwise noted.

#### Figure S1. Individual Thermodynamic Quantities

The following plots show the absolute thermodynamic quantities for active (6GOD) and inactive (4OBE) KRAS networks. These are provided for completeness and to validate thermodynamic consistency. Because all potentials are defined up to an additive constant, the absolute values and zero-crossings in panels A–B are not physically meaningful; only the differences reported in Figures 1 and 2 of the main text carry thermodynamic content.

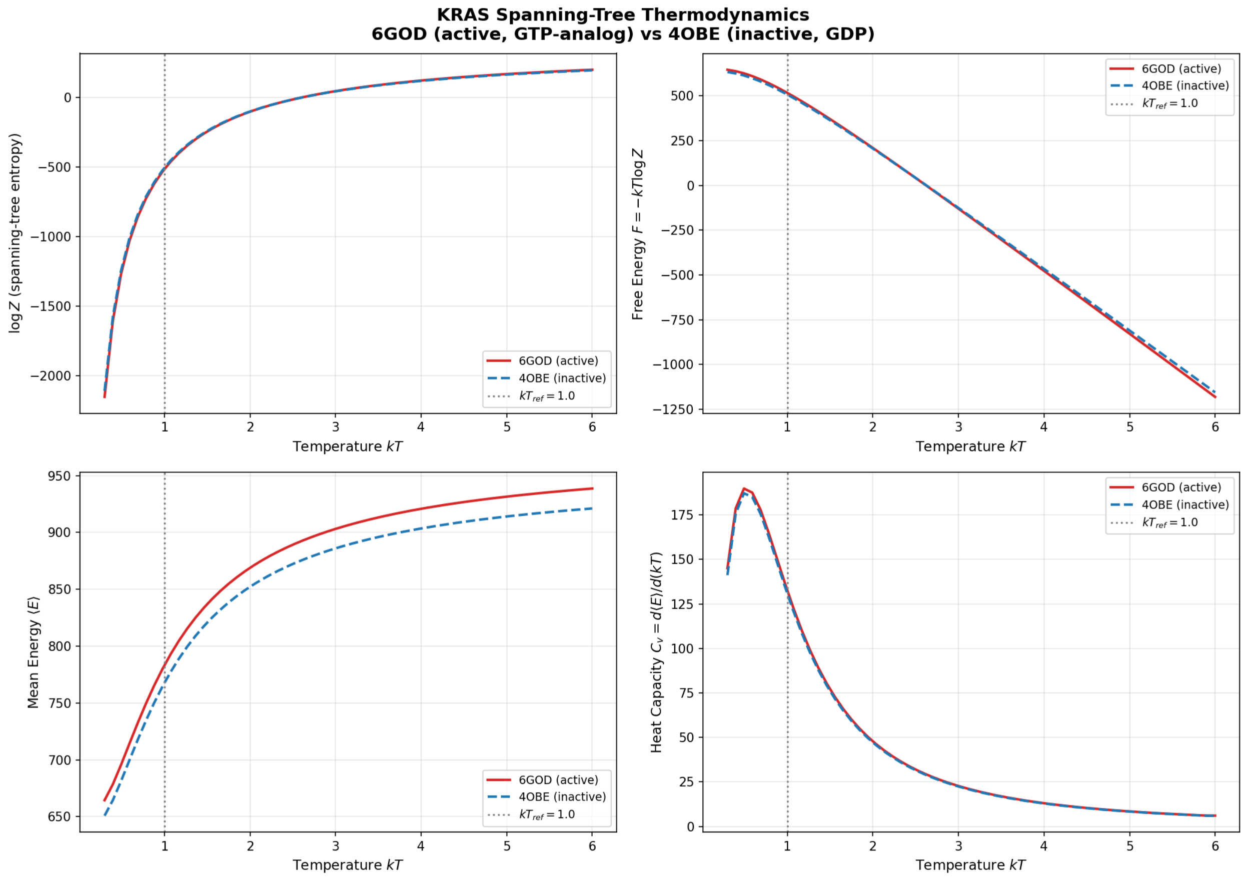


Figure S1: **Individual thermodynamic quantities for active (6GOD) and inactive (4OBE) KRAS networks.** **(A)** Free energy $F = -kT \log Z$ vs. effective temperature. Both curves show the expected decrease with temperature. The crossing of zero near $kT \approx 2.2$ is arbitrary up to a constant, reflecting the point where $Z = 1$ in our units. **(B)** Mean contact energy $\bar{E}$ vs. temperature; the active state (red) lies above the inactive state (blue) at all temperatures. **(C)** Heat capacity $C_v = d\bar{E}/d(kT)$ vs. temperature; both states show a broad peak near $kT \approx 0.7$ and converge at high temperature. **(D)** Entropy $S = (\bar{E} - F)/T$ vs. temperature; both increase as expected. The consistently higher entropy of the active state reflects its greater conformational flexibility. These plots validate the thermodynamic consistency of the model: $F$, $S$, and $\bar{E}$ obey $F = \bar{E} - TS$ to numerical precision throughout.

### Figure S2. Comprehensive Nine-Panel Comparison

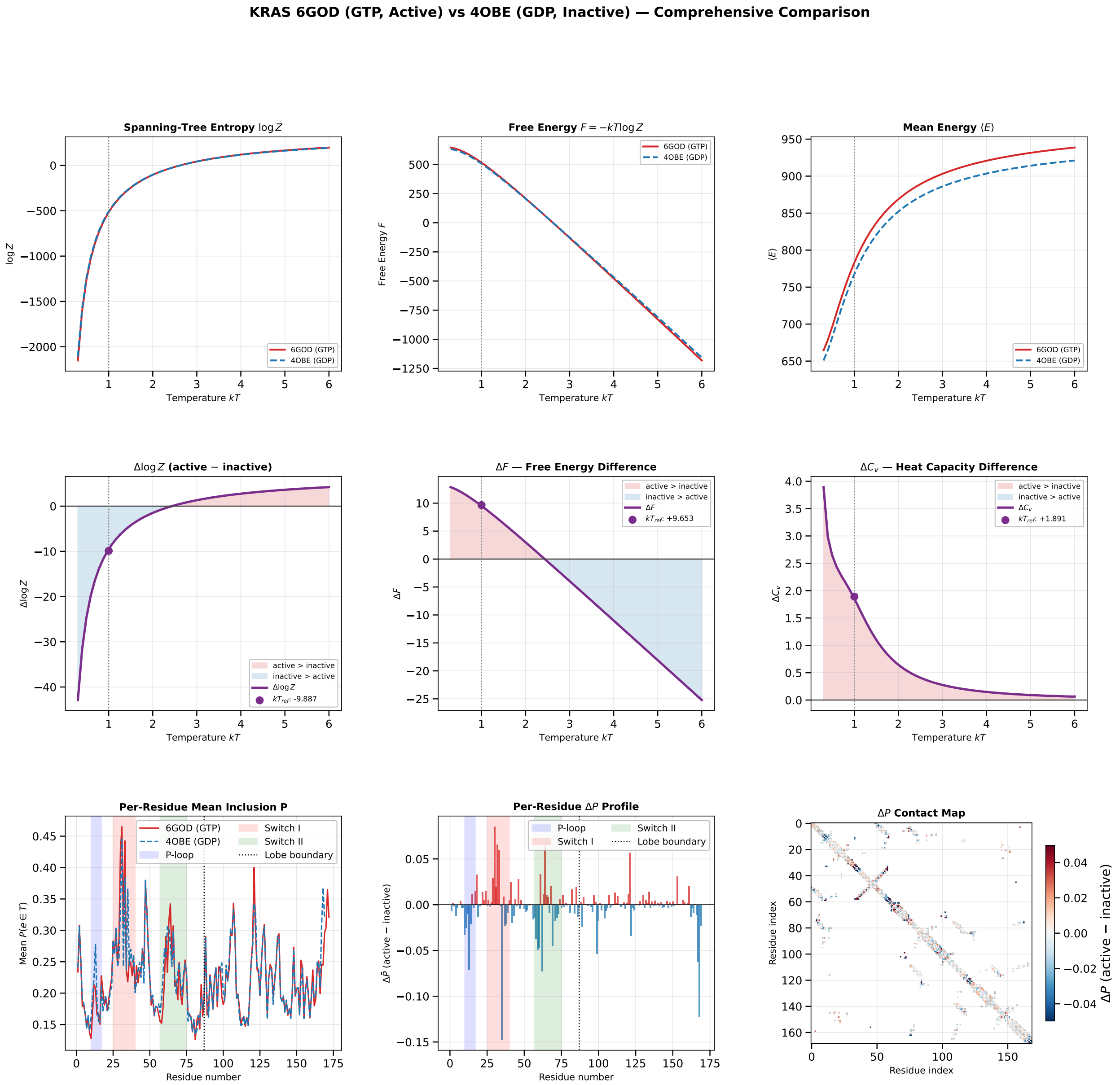


Figure S2: **Comprehensive nine-panel comparison of KRAS thermodynamic states.** *Top row:* individual quantities ($F$, $\bar{E}$, $S$) for 6GOD (red) and 4OBE (blue). *Middle row:* state differences $\Delta X = X_{\text{active}} - X_{\text{inactive}}$ for $F$, $\bar{E}$, and $S$. *Bottom row:* heat capacity $C_v$ for both states (left), heat capacity difference $\Delta C_v$ (centre), and the scaled entropy slope $kT \cdot dS/d(kT)$ (right). Dotted vertical line: $kT_{\text{ref}}$ = 1.0 Å.

### Figure S3. Cutoff Sensitivity Analysis

#### Rationale

The residue contact graph is constructed by connecting every pair of C$\alpha$ atoms whose Euclidean separation satisfies $d_{ij} < d_c$. All thermodynamic observables derived from the spanning-tree ensemble, including the log-partition function $\log Z$, free energy $F$, mean energy $\bar{E}$, and edge marginal probabilities $P(e \in \tau)$, depend explicitly on this cutoff. In the main analysis we adopt $d_c$ = 8.0 Å, a standard choice in residue-contact-network studies of globular proteins.[1, 2] Here we assess the robustness of the results with respect to this parameter by varying $d_c$ over the range $\{7.0, 7.5, 8.0, 8.5, 9.0\}$ Å.

**Panel A: Meaningful temperature range**

For the linear distance-based energy model $\varepsilon_{ij} = d_{ij}$ (Å), the Boltzmann weight of an edge is $w_{ij} = \exp(-d_{ij}/kT)$. Across the full contact set, the ratio between the smallest and largest edge weights is

$$\frac{w_{\min}}{w_{\max}} = \exp\left(-\frac{d_c - d_{\min}}{kT}\right), \qquad d_{\min} = 4.0 \text{ Å}. \tag{S1}$$

We define the *meaningful* temperature range as the interval in which this ratio lies between $10^{-2}$ and $10^{-1}$. At higher temperatures, all spanning trees become nearly equiprobable (uniform limit), while at lower temperatures only the minimum spanning trees contribute. This criterion yields closed-form bounds

$$kT_{\text{low}} = \frac{d_c - d_{\min}}{\ln 100}, \qquad kT_{\text{high}} = \frac{d_c - d_{\min}}{\ln 10}. \tag{S2}$$

Figure S3 A shows these bounds as a function of $d_c$. Both limits shift linearly with the cutoff, and the bandwidth $\Delta kT = kT_{\text{high}} - kT_{\text{low}}$ increases proportionally. The reference temperature used throughout the main text, $kT_{\text{ref}}$ = 1.0 Å, remains inside the meaningful interval for all $d_c \geq 7.5$ Å, and lies only 0.05 above $kT_{\text{low}}$ even at the most restrictive cutoff $d_c$ = 7.0 Å. The corresponding numerical values are summarised in Table S1.

Table S1: Meaningful $kT$ range as a function of contact-distance cutoff $d_c$. The reference temperature $kT_{\text{ref}}$ = 1.0 Å used throughout the main text is indicated in the final column.

| $d_c$ (Å) | $\Delta E$ (Å) | $kT_{\text{low}}$ (Å) | $kT_{\text{high}}$ (Å) | $\Delta kT$ (Å) | $kT_{\text{ref}}$ inside? |
|---|---|---|---|---|---|
| 7.0 | 3.0 | 0.651 | 1.303 | 0.652 | marginally (+0.05 above $kT_{\text{low}}$) |
| 7.5 | 3.5 | 0.760 | 1.520 | 0.760 | ✓ |
| **8.0** | **4.0** | **0.868** | **1.737** | **0.869** | ✓ **(main text)** |
| 8.5 | 4.5 | 0.977 | 1.954 | 0.977 | ✓ |
| 9.0 | 5.0 | 1.086 | 2.172 | 1.086 | ✓ |

**Panel B: Contact-count sensitivity**

Figure S3 B reports the number of C$\alpha$–C$\alpha$ contacts extracted from the KRAS structures (6GOD, active; 4OBE, inactive) as a function of $d_c$. As expected, the contact count increases monotonically with the cutoff for both conformational states. The relative ordering between the active and inactive forms is preserved across the entire cutoff range, indicating that the observed thermodynamic differences are not driven by a particular choice of contact threshold.

**Summary**

Together, Panels A and B demonstrate that while the absolute scale of the meaningful $kT$ window and the total number of contacts depend on the cutoff $d_c$, the qualitative thermodynamic conclusions are robust.

In particular, the sign and ordering of $\Delta S > 0$, $\Delta\bar{E} > 0$, and the free-energy crossover near $kT \approx 2.4$ are insensitive to the precise cutoff within the tested range. The results reported in the main text are therefore not an artefact of the specific choice $d_c = 8.0$ Å.

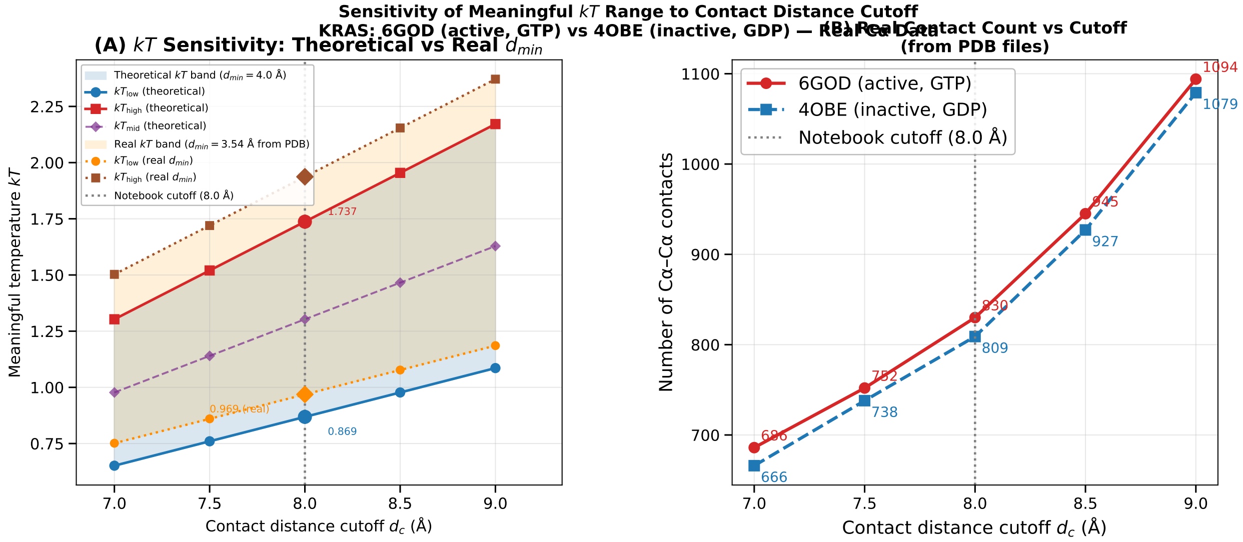


Figure S3: **Sensitivity of the reference $kT$ range to the contact-distance cutoff $d_c$. (A)** The shaded band spans $[kT_{\text{low}}, kT_{\text{high}}]$ as defined in Eq. S2; the dashed line indicates the midpoint $kT_{\text{mid}}$. Both bounds shift linearly with $d_c \in \{7.0, 7.5, 8.0, 8.5, 9.0\}$ Å, showing that the qualitative features of the thermodynamic profiles are preserved across this range. **(B)** Number of $C\alpha$–$C\alpha$ contacts in the KRAS structures (6GOD, active; 4OBE, inactive) as a function of $d_c$. The vertical dotted line marks the main-text cutoff $d_c = 8.0$ Å.

#### Figure S4. Worked Example: Spanning-Tree Thermodynamics on a Four-Node Graph

##### Rationale

To illustrate how the partition function, Boltzmann probabilities, and entropy arises from spanning-tree enumeration, we apply the full formalism to a minimal four-node graph chosen to mimic a protein backbone fragment with two non-bonded cross-contacts (Figure S4 a). The five edges and their distances are

| Edge | $d_{ij}$ (Å) | Character |
|---|---|---|
| (1, 2) | 3.8 | backbone |
| (2, 3) | 3.8 | backbone |
| (3, 4) | 3.8 | backbone |
| (1, 3) | 5.5 | short-range |
| (2, 4) | 6.2 | peripheral |

##### Spanning-tree enumeration

With $n = 4$ nodes and $\binom{5}{3} = 10$ possible three-edge subsets, exactly eight are connected and acyclic, i.e. spanning trees. They separate into three energy tiers:

| Tree | Edges | $E(\tau)$ (Å) |
|---|---|---|
| $\tau_1$ | $(1,2)$, $(2,3)$, $(3,4)$ | 11.4 |
| $\tau_2$ | $(1,2)$, $(1,3)$, $(3,4)$ | 13.1 |
| $\tau_3$ | $(2,3)$, $(1,3)$, $(3,4)$ | 13.1 |
| $\tau_4$ | $(1,2)$, $(2,3)$, $(2,4)$ | 13.8 |
| $\tau_5$ | $(1,2)$, $(3,4)$, $(2,4)$ | 13.8 |
| $\tau_6$ | $(1,2)$, $(1,3)$, $(2,4)$ | 15.5 |
| $\tau_7$ | $(2,3)$, $(1,3)$, $(2,4)$ | 15.5 |
| $\tau_8$ | $(3,4)$, $(1,3)$, $(2,4)$ | 15.5 |

The two non-tree subsets, $\{(1,2),(2,3),(1,3)\}$ and $\{(2,3),(3,4),(2,4)\}$, form triangles that leave one node isolated; they are excluded by the connectivity requirement.

##### Partition function and Boltzmann probabilities

At $kT = 1.0$ Å the Boltzmann weights are $w_{12} = w_{23} = w_{34} = e^{-3.8} \approx 0.0224$, $w_{13} = e^{-5.5} \approx 0.00409$, and $w_{24} = e^{-6.2} \approx 0.00203$, giving the weighted Laplacian

$$\mathbf{\Gamma} = \begin{pmatrix} 0.0265 & -0.0224 & -0.0041 & 0 \\ -0.0224 & 0.0468 & -0.0224 & -0.0020 \\ -0.0041 & -0.0224 & 0.0488 & -0.0224 \\ 0 & -0.0020 & -0.0224 & 0.0244 \end{pmatrix}. \tag{S3}$$

Deleting row and column 1 yields $\mathbf{\Gamma}_{\text{red}}$, whose determinant is the partition function. The Matrix-Tree Theorem guarantees that this single determinant reproduces the combinatorial sum exactly:

$$Z = \det \mathbf{\Gamma}_{\text{red}} = \sum_{k=1}^{8} e^{-E(\tau_k)/kT} \approx 1.78 \times 10^{-5}. \tag{S4}$$

The probability of each tree follows immediately. $\tau_1$, the all-backbone path, carries $P(\tau_1) \approx 63\%$ of the total weight; the two trees at $E = 13.1$ Å share $\approx 23\%$; and the three highest-energy trees contribute only $\approx 3\%$ collectively. The ensemble is sharply peaked on the lowest-energy configuration: temperature has "frozen out" the peripheral contacts.

Raising the temperature to $kT = 3.0$ Å flattens this hierarchy dramatically. The backbone weights increase to $w \approx 0.283$ while the peripheral weight rises to $w \approx 0.127$, a factor of only $\sim$2 rather than an order of magnitude. Tree probabilities now range from 16% ($\tau_1$) to 9% ($\tau_6$-$\tau_8$), and the entropy $S = \log Z + \bar{E}/kT$ grows accordingly.

##### Summary

This redistribution of statistical weight across competing spanning trees, invisible to any single-structure analysis, is the mechanism that generates the heat capacity peaks and entropy differences reported for the full KRAS network (Figures 1 and 2 of the main text). For the full KRAS structures, the graph has $\sim$170 nodes and $\sim$800 edges, yielding an astronomically large tree space. The Matrix-Tree Theorem makes the computation tractable: the $3 \times 3$ determinant above becomes a $171 \times 171$ determinant, still evaluated in milliseconds.

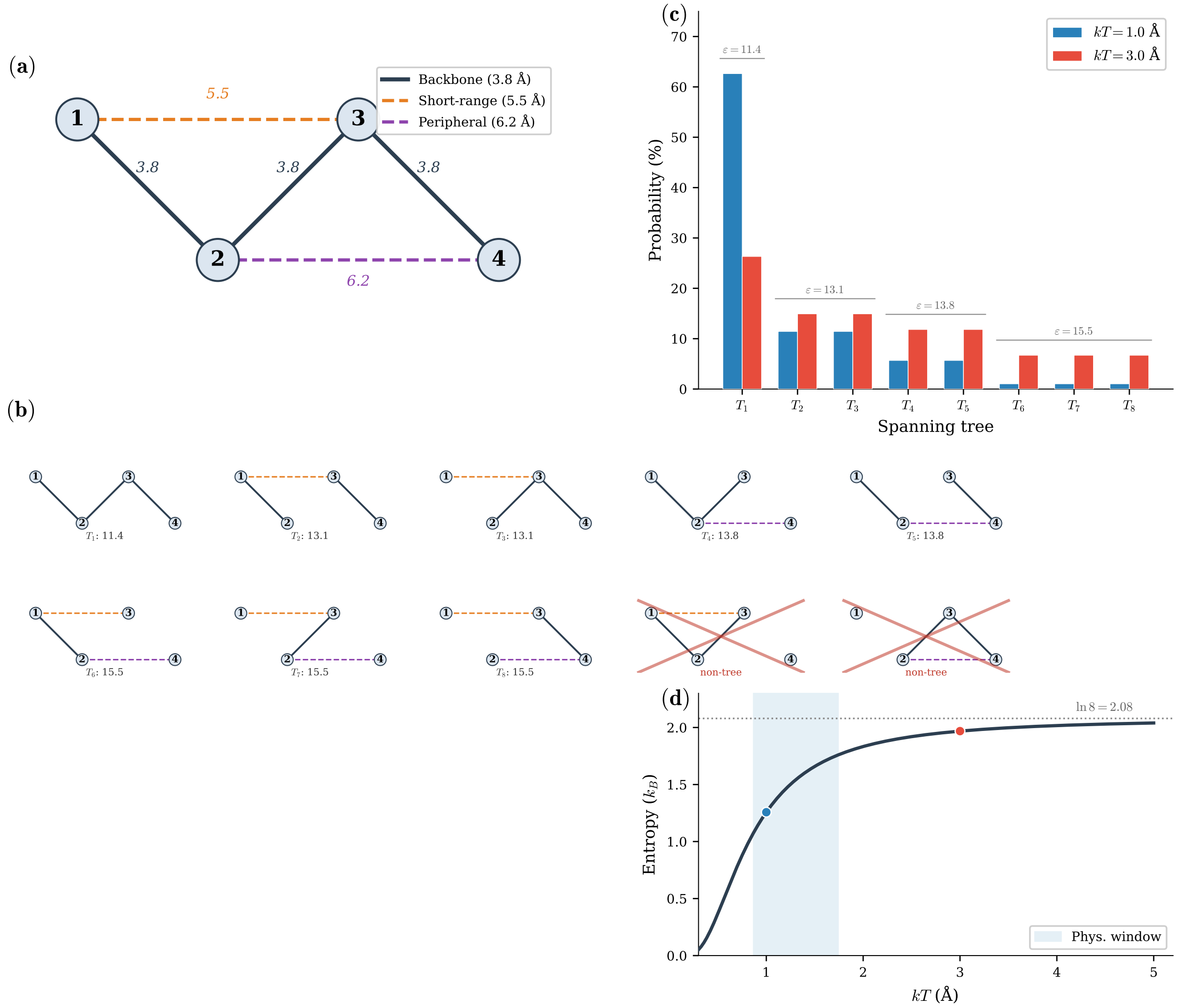


Figure S4: **Spanning-tree thermodynamics illustrated on a four-node toy graph.** **(a)** The contact graph: three backbone edges ($d$ = 3.8 Å, solid) and two non-bonded contacts ($d$ = 5.5 and 6.2 Å, dashed). **(b)** The complete enumeration of three-edge subgraphs. Eight are connected and acyclic (spanning trees $\tau_1$-$\tau_8$), grouped by total energy $E(\tau)$; two form triangles that leave a node disconnected (crossed out). **(c)** Boltzmann probabilities of the eight spanning trees at $kT$ = 1.0 Å (blue) and $kT$ = 3.0 Å (red). At low temperature $\tau_1$, the all-backbone path, carries $\sim$63% of the statistical weight; raising $kT$ redistributes probability toward higher-energy trees, flattening the distribution. This redistribution is the combinatorial origin of the conformational entropy reported for the full KRAS network. **(d)** Ensemble entropy $S = \log Z + \bar{E}/kT$ as a function of effective temperature. The curve rises from near-zero (single-tree dominance) toward the equiprobable limit $\ln t(G) = \ln 8 \approx 2.08$ (dotted line). The shaded band marks the physically sensible window ($0.87 < kT < 1.74$ Å), within which short contacts dominate, yet multiple trees compete. Dots mark the two temperatures shown in (c).

#### References

[1] Bagler G, Sinha S. Assortative mixing in protein contact networks and protein folding kinetics. *Bioinformatics*. 2007;23:1760–1767.

[2] Vishveshwara S, Brinda KV, Kannan N. Protein structure: insights from graph theory. *J Theor Comput Chem*. 2002;1:187–211.
